# MOFA: Multi-Objective Flux Analysis for the COBRA Toolbox

**DOI:** 10.1101/2021.05.20.445041

**Authors:** Marc Griesemer, Ali Navid

## Abstract

Multi-objective Optimization (MO) is an important tool for quantitative examination of the trade-offs faced by biological organisms. Using genome-scale constraint-based models of metabolism (GSMs), Multi-Objective Flux Analysis (MOFA) allows MO analyses of trade-offs among key biological tasks. The leading software package for conducting a plethora of different types of constraint-based analyses using GSMs is the COBRA Toolbox for MATLAB. We have developed a new add-on tool for this toolbox using Normalized Normal Constraint (NNC) that performs MOFA for a number of objectives only limited by computation power (*n* ≤ 10). This development will facilitate MOFA analyses by COBRA’s large user base and allow greater multi-faceted examination of metabolic trade-offs in complicated biological systems.

**Availability and Implementation:** The MOFA software is freely available for download from https://bbs.llnl.gov under the GPL v2 license. The program runs on MATLAB with the COBRA software on Windows, Linux, and MacOS. It includes a detailed manual explaining the input and output of a simulation, a listing of the code’s functions, and an example MOFA run using a well-curated GSM model of *E. coli*.

## 1 Introduction

Evolution necessitates that organisms maximize in a Pareto-optimal fashion a myriad of different biological objectives to optimize an organism’s fitness under different environmental conditions. A Pareto-optimal outcome among biological tasks is one where improvement in performance of one task would result in diminishment of the ability to achieve one or more other tasks. Sufficient knowledge about biological systems to manipulate their workings requires quantitative understanding of these tradeoffs. GSMs are a key tool for studying biological systems and constraint-based methods such as Flux Balance Analysis (FBA) (Orth *et al*., 2010) are the most popular mathematical methods for assessing system characteristics. The Constraint-Based Reconstruction and Analysis (COBRA) Toolbox (Schellenberger *et al*., 2011) is a popular open-source code base addition to MATLAB (Natick, MA) for simulating FBA and other constraint-based system-level analyses.

When using typical FBA, linear programming is used to solve for a feasible biochemical flux pattern that would result in optimum operation of one biological objective, typically growth, at steady state conditions. However, to assess tradeoffs between multiple biological objectives (e.g., growth, regulatory, or defense related processes in a single cell; or optimum growth for different cell types in a community) one needs to conduct a Multi-Objective Flux Analysis (MOFA) of the system. MOFA maps the *n*-dimensional (*n*=number of examined objectives) surface of the Pareto front by calculating a large set of Pareto solutions. However, calculating the solution for a relatively large number of objectives (e.g., *n* ≥ 10) can be computationally costly and require days or weeks of simulation time on a typical desktop computer. To our knowledge, currently the COBRA toolbox lacks a stand-alone module to conduct MOFA analyses for more than three objectives. That capability is not sufficient for studying tradeoffs among an organisms numerous bioproducts and growth. In addition, individual species in microbial communities often have conflicting goals for growth with each needing to be a separate objective in a simulation. Here we report development of a new MATLAB code that can be used with the COBRA toolbox to conduct MOFA analyses for as many objectives as processing power allows (currently *n* ≤ 20). Our motivations for developing this code were to: 1) allow for facile examination of metabolic tradeoffs using GSMs in COBRA, 2) permit simultaneous examination of a high (relative to most previous systems biology studies) number of objectives, and 3) evenly map the Pareto front regardless of the magnitudes of the examined objectives.

## 2 Features and Implementation

### 2.1 Simulation Algorithm

The code uses the Normalized Normal Constraint (NNC) (Messac *et al*., 2003) method for MO analyses to ensure a full set of evenly distributed Pareto optimal points regardless of the differences in magnitudes of the examined objectives. A description of the NNC is given in the original paper as well as in the provided user manual.

### 2.2 Software Implementation

Our software initializes with a MATLAB COBRA model object containing the user’s appropriately constrained GSM. The user can import the model into this format from a SBML file and then make any needed modifications to the object structure such as changes in constraints before calling the MOFA function. The code uses the open-source GLPK linear programming solver installed with COBRA. The size of the problem is proportional to the number of objectives being examined. It is also dependent on how many Pareto solutions are used to map the Pareto front. The number of divisions (*x*) along the normalized length of each objective can significantly impact the time needed to conduct MOFA analyses. Another factor that determines the run time for a simulation is the relationship between the examined objectives. If two or more objectives do not negatively affect the value of the others, then the problem will be less complicated and the simulation will finish sooner. Results and run times may vary, even among different sets of objectives within the same model.

Our MOFA algorithm is practical for calculating the Pareto front of a MO problems with up to 10 objectives on a laptop computer in one day. For the results shown in Figure 1, we used an Intel Xeon-based workstation and a personal laptop. Using the desktop machine, MOFA analysis (*n* = 10, *x* = 10) using our code on a well-curated GSM of *E. coli*, iAF1260 (Feist *et al*., 2007), took 3.0 hours. On the laptop, it took 26.1 hours. With *n* ≫ 10 objectives, the use of high-performance computing (HPC) is necessary and beyond the aims of our MATLAB MOFA software. As viewed in Figure 1, the simulation time does not scale linearly with the number of objectives.

**Figure 1:**
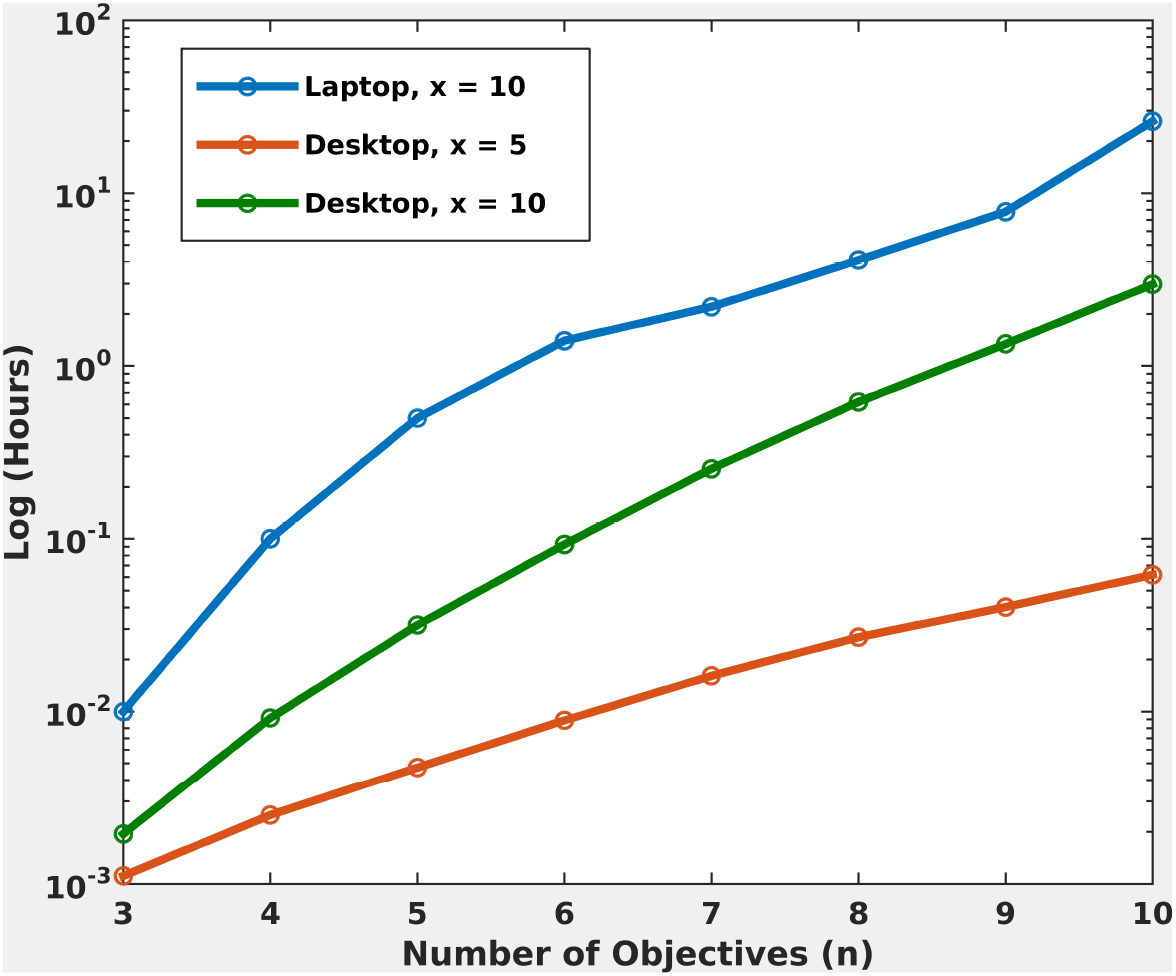
Plot illustrating the run time of MOFA for the E. coli model iAF1260 [4] on both a desktop workstation (Intel Xeon E5-2697 v3 (4 core), 3.5 GHz, 32 GB, Linux Red Hat) and personal laptop (Intel Core i7-2720QM (2-core), 2.2 GHz, 8 GB, Windows 7). The number of divisions (*x*) gives the granularity of the space intervals; however, 10 divisions take approximately 35 times longer than 5 (using 10 objectives).

Throughout our algorithm, we have devised various checks and short cuts to speed up the simulation time while maintaining fidelity of our results. For example, we speed up the algorithm by avoiding calculation of new Pareto optimal solutions in regions of the solution space that are very close to previously determined solutions.

## 3 Conclusion

In conclusion, our code is an addition to the COBRA toolbox that facilitates the use of the MOFA technique for a large user community. It provides a solution in a timely manner and allows for efficient analysis of costs and tradeoffs for a relatively large number of objectives.

## Supporting information

Operation manual

## Acknowledgements

Work was performed under the auspices of the U.S. Department of Energy by Lawrence Livermore National Laboratory under Contract DE-AC52-07NA27344. We would like to thank Peter Weber and Dan Kirshner for their editing of the manuscript and their advice.

