## Supplementary material for "MOFA: Multi-Objective Flux Analysis for the COBRA Toolbox": Operation manual

### MOFA User Manual

Marc Griesemer    Ali Navid

**5 An example MOFA analysis using an *E. coli* model** **6**

**6 Normalized Normal Constraint (NNC) method** **9**

**7 Definitions** **12**

**References** **13**

### 1 Introduction to MOFA

MOFA is a COBRA toolbox (Schellenberger *et al.*, 2011) extension using MATLAB for performing multi-objective analysis of trade-offs among different objectives of a biosystem using constraint-based models of biochemical processes. This document outlines the software. Any queries regarding use of MOFA are welcome and should be directed to Marc Griesemer or Ali Navid.

### 2 User Requirements

#### 2.1 Hardware and Software Requirements

No specific hardware requirements are necessary beyond what MATLAB needs. The code uses the GLPK solver that is included with the latest download of COBRA Toolbox. No other LP solvers are currently supported.

#### 2.2 Simulation Requirements

##### 2.2.1 User Considerations

1. The user needs to ensure that model is well constrained and provides accurate FBA solutions that do not violate mass balance and thermodynamic laws.

2. The user must confirm that the model would be valid (without errors) for optimization using COBRA toolbox.

##### 2.2.2 Structure of Input Arguments

MOFA needs information on the parameters of the simulation: namely, the test model, the number of divisions of the Pareto frontier, and the specified objectives. The user can use an input file (see Table M1) or function arguments to supply the list of objectives. The program uses a modified constraint-based COBRA model object that has unique constraints for the defined objectives. The solution is the $n$-dimensional ($n$=number of examined objectives) solution of the normalized objective values that composes the Pareto front. One must use valid objectives contained in the model; otherwise, the program will print an error message and exit. It is useful to check

Table M1: Structure of input file includes list of objectives, including the main objective.

| (name of file).txt |
| --- |
| # COMMENT<br>(name of the objective to be optimized during MOFA iterations)<br>#COMMENT<br>(objective name 1)<br>(objective name 2)<br>...<br>(objective name n) |

to that the objective names are correctly spelled and that there are no blank lines at the end of the list.

##### 2.2.3 Runtime Factors

The time taken by the simulation depends on:

1. the number of objectives
2. the number of divisions
3. the objectives chosen

### 3 Installation

1. Install MATLAB (`http://www.mathworks.com`); a license is required.
2. Download and install the COBRA toolbox as directed [1].
3. Install MOFA by adding the package folder into the COBRA folder. Alternative locations outside the MATLAB path for MOFA will require a path addition.

### 4 MOFA Function Structure

#### 4.1 Index of MOFA functions

| | | |
|---|---|---|
| 1 | mofa.m | main function |
| 2 | min_max.m | finds the minimum and maximum fluxes of all objectives |
| 3 | anch_pts.m | finds the anchor points, auxilary function |

#### 4.2 Functions for running a simulation

There are the two functions the user can use to conduct MOFA analyses, *mofa* and *min_max*. A third function, *anch_pts*, determines the anchor points but is an auxiliary function used only by the main MOFA program. This subsection gives a listing of the inputs and outputs of these functions.

##### 4.2.1 mofa.m

This is the main MOFA function for the code.

function [mofa_sol, mxhr, mihr, aphr] = ***mofa***(model, inp_file, obj, obf, ndiv, mi_mx)

Inputs:

| | |
|---|---|
| model | COBRA model object |
| inp_file | Input text filename with listing of objectives (string) |
| obj | Optional: list of objectives (cell array of strings).<br>Alternatively, put in inp_file. |
| obf | Optional: main objective (1 cell string).<br>Alternatively, put in inp_file. |
| ndiv | Optional: number of divisions (integer) |
| mi_mx | Optional: Main objective minimized or maximized<br>(string, 'min' or 'max') |

Outputs:

| | |
|---|---|
| mofa_sol | List of feasible values for Pareto points (matrix of doubles) |
| mxhr | $N$ member array (doubles) containing the calculated maximum values of each objective. |
| mihr | $N$ member array (doubles) containing the calculated minimum values of each objective. |
| aphr | 2-D matrix (doubles) containing the anchor points. Its size is $(N \times N)$, where $N$ is the number of objectives. |

##### 4.2.2 min_max.m

This function is used by the main MOFA function but can be called independently.

function [mxhr,mihr] = ***min_max***(model, objc)

This function solves for the maximum and minimum fluxes of each objective. This is another way of conducting flux variability analysis (FVA) (Mahadevan and Schilling, 2002). FVA is a method in COBRA that solves for the upper and lower bounds of all steady-state reaction fluxes in a model. In the MOFA code, this is done only for the objectives of interest.

Inputs:

| | |
|---|---|
| model | COBRA model object |
| objc | the names of the objectives (cell array of strings) |

Outputs:

| | |
|---|---|
| mxhr | $N$ member array (doubles) containing the calculated maximum values of each objective. |
| mihr | $N$ member array (doubles) containing the calculated minimum values of each objective. |

### 5 An example MOFA analysis using an *E. coli* model

First, initiate the COBRA toolbox:

```
>> initCobraToolbox
```

GLPK is the only supported solver at this time. If it is not the current solver, then the program will switch to it for MOFA usage.

The *E. coli* model (iAF1260 (Feist *et al.*, 2007)) is contained in an SBML file ('Ec_iAF1260_flux1.xml') at the URL:https://www.ncbi.nlm.nih.gov/pmc/articles/PMC1911197/ (Supplementary Information 6 at reference). We chose iAF1260 as an example because it is one of the best currently available human-curated models. The example MOFA analyses are meant solely to demonstrate the workings of our MOFA code, rather than to provide novel biological insight. The model should be imported to a COBRA model object.

The model can be loaded into MATLAB using the following command:

```
>> model = readCbModel('Ec_iAF1260_flux1.xml');
```

Then navigate to the directory where the MOFA folder is located.
There are two different ways to input the objectives into the program:

1. Typing in the objectives into variables *obj* and *obf*:

   ```
   >> obj = { 'EX_o2(e)', 'EX_co2(e)', 'EX_ac(e)', 'EX_etoh(e)',
   'EX_nh4(e)'};
   ```

   These represent exchange reactions for oxygen, carbon dioxide, acetate, ethanol, and ammonia, respectively.

   The main objective (objective to be optimized) also needs to be specified:

   ```
   >> obf = {'Ec_biomass_iAF1260_core_59p81M'};
   ```

2. Using an input file:

   One can also use an input file to read in the names of the objectives in which case the third and fourth arguments must be empty. The input file for the E. coli simulation with 6 objectives is shown in Table M2.

Table M2: Input file for the *E. coli* with objectives chosen from the model's reactions ('mofa_ecoli_input.txt').

```
#enter the name of the objective to be optimized
Ec_biomass_iAF1260_core_59p81M
#enter the list of other objectives
EX_o2(e)
EX_co2(e)
EX_ac(e)
EX_etoh(e)
EX_nh4(e)
```

For a full list of available reactions, examine the 'rxns' and 'rxnNames' fields in the COBRA model object.

Finally, the number of divisions can also be specified:

**>> ndiv = 10;**

(The number of divisions must be positive and greater than 3.)
At this point, one can look at the model object and make modifications to the constraints and other fields as necessary before calling the main MOFA function.

Depending on how the objectives were specified, the arguments to the main function are handled differently. If they were placed in cell arrays of strings, then the MOFA function can look like:

**>> [mofa_sol, mxhr, mihr, aphr] = mofa(model, [] , obj, obf, ndiv, 'max');**

In this case, no input file is used to import the list of objectives, so the second argument is empty.

If the input file was used, then the second argument contains the name of the input file and the third and fourth arguments for the objective arrays are empty.

**>> [mofa_sol, mxhr, mihr, aphr] = mofa(model,'mofa_ecoli_input.txt' , [] , [] , ndiv, 'max');**

To review, the first argument is the model as a COBRA object; the second is the name of an input file containing the list of objectives; the third is the list of objectives; the fourth gives the main objective; and the fifth and sixth are the number of divisions and whether to maximize ('max') or minimize ('min') the main objective, respectively. Only the first two arguments are mandatory in which case an input file is necessary. The last four are optional. If the number of divisions and optimization sense are not supplied, the number of divisions is 10 and the main objective is maximized.

Table M3: Sample output of the *E. coli* simulation illustrating the values of the individual objectives at each point for the feasible solutions of the Pareto front (see also 'mofa_output.txt').

| EX_ac(e) | EX_co2(e) | EX_etoh(e) | EX_nh4(e) | EX_o2(e) | Biomass |
|---|---|---|---|---|---|
| 0.30 | 0.22 | 0.30 | 0.00 | 0.13 | 0.00 |
| 0.00 | 0.37 | 0.00 | 0.22 | 0.88 | 1.00 |
| 1.00 | 0.06 | 0.00 | 0.00 | 0.15 | 0.00 |
| 0.00 | 0.62 | 0.10 | 0.11 | 0.79 | 0.50 |
| 0.60 | 0.43 | 0.00 | 0.00 | 0.44 | 0.00 |
| 0.10 | 0.72 | 0.00 | 0.07 | 0.81 | 0.30 |
| 0.10 | 0.40 | 0.20 | 0.13 | 0.63 | 0.60 |
| 0.40 | 0.29 | 0.50 | 0.00 | 0.15 | 0.00 |
| 0.10 | 0.32 | 0.20 | 0.09 | 0.46 | 0.40 |
| 0.60 | 0.15 | 0.00 | 0.07 | 0.35 | 0.30 |
| 0.80 | 0.12 | 0.10 | 0.02 | 0.21 | 0.10 |
| 0.00 | 0.40 | 0.00 | 0.18 | 0.79 | 0.80 |

The output of the MOFA code is a table of columns representing objectives and rows representing a Pareto solution in the n-dimensional objective space. The results are normalized, i.e., the fraction of the optimum value that the objective can attain given the model constraints. Thus, the values will be between 0 and 1. All of the output values are provided both in a tab-separated data file (Table M3) and in MATLAB variables for further analysis and graph plotting using MATLAB or Excel.

### 6 Normalized Normal Constraint (NNC) method

Our MOFA code uses the NNC method of multi-objective optimization (Messac *et al.*, 2003).

**Problem Statement:** The multi-objective optimization (MO) problem can be defined as:

$$\min_x \{F_1(x)\ F_2(x)\ \cdots\ F_n(x)\},\ n \geq 2 \tag{1}$$

subject to the constraints:

$$g_j(x) \leq 0, 1 \leq j \leq r \tag{2}$$

$$h_k(x) = 0, 1 \leq k \leq s \tag{3}$$

$$x_{li} \leq x_i \leq x_{ui},\ \ 1 \leq i \leq n_x \tag{4}$$

The vector $x$ denotes the set of constraint variables and $F_i$ denotes the $ith$ objective.

Algorithm Steps:

**Step 1: Initialize Simulation**

a. Load SBML model or COBRA model object.
b. Process input file or lists of objectives.

**Step 2: Find utopia/nadir points (maximum/minimum flux values for all objectives).**

The points that contain the set of maximum and minimum values of all objectives are called the

1. Utopia Point
$$F^U = [F_1(x^{1*})\ F_2(x^{2*}) \cdots F_n(x^{n*})] \tag{5}$$

2. Nadir Point
$$F^N = [F_1^N\ F_2^N \cdots F_n^N] \tag{6}$$

where $F_i^N = \max[F_i(x^{1*})\ \ F_i(x^{2*}) \cdots F_i(x^{n*})]$, $i \in \{1, 2, \ldots, n\}$.

These points are used as a reference points in objective space as there is not a way to simultaneously optimize all objectives. In the NNC, the utopia point is used to normalize the space in each dimension. This is essentially performing Flux Variability Analysis (FVA) on the defined objectives.

The difference between these two points gives the range for each dimension in objective space, a vector:

$$\bar{F} = \left\{ \begin{array}{c} \nu_1 \\ \nu_2 \\ \vdots \\ \nu_n \end{array} \right\} = F^N - F^U \tag{7}$$

which leads to the normalized objectives,

$$\bar{F}_i = \frac{F_i - F_i(x^{i*})}{\nu_i},\ i \in \{1, 2, \ldots, n\} \tag{8}$$

**Step 3: Find Anchor Points**

The NNC method uses anchor points as reference vertices in objective space. The anchor points are calculated by individually minimizing each objective ($F_j$) individually, subject to the problem constraints, to obtain the *jth* anchor point, $F^{j*}$ $(j = 1, ..., n)$.

**Step 4: Define utopia line (utopia hyperplane).**

From the vertices of the anchor points, we can define a utopia hyperplane. We define the direction of the utopia line vector

$$\bar{N} = \bar{F}^{n*} - \bar{F}^{k*} \tag{9}$$

Then we compute a normalized increment along the direction $\bar{N}_k$ for a prescribed number of divisions $x$ for each direction $k$:

$$\delta_k = \frac{1}{x-1}, 1 \leq k \leq n-1 \tag{10}$$

Parameter $\alpha_{ij}$ is incremented by $\delta$ between 0 and 1 and we use values of $j$ where $j \in \{1, 2, \ldots, n\}$.

**Step 5: Generate evenly distributed hyperplane points.**

Evaluate a set of evenly distributed points on the Utopia hyperplane as

$$X_{pj} = \sum_{j=1}^{n} \alpha_{kj} \bar{F}^{k*} \tag{11}$$

where

$$0 \leq \alpha_{kj} \leq 1 \tag{12}$$

and

$$\sum_{k=1}^{n} \alpha_{kj} = 1. \tag{13}$$

Next the use the set of equally distributed points generated at the previous step to compute the Pareto solution by solving the following LP problem at each point individually

$$\min \bar{F}_n \tag{14}$$

$$g_j(x) \leq 0, 1 \leq j \leq r \tag{15}$$

$$h_k(x) = 0, 1 \leq k \leq s \tag{16}$$

$$x_{li} \leq x_i \leq x_{ui},\ \ 1 \leq i \leq n_x \tag{17}$$

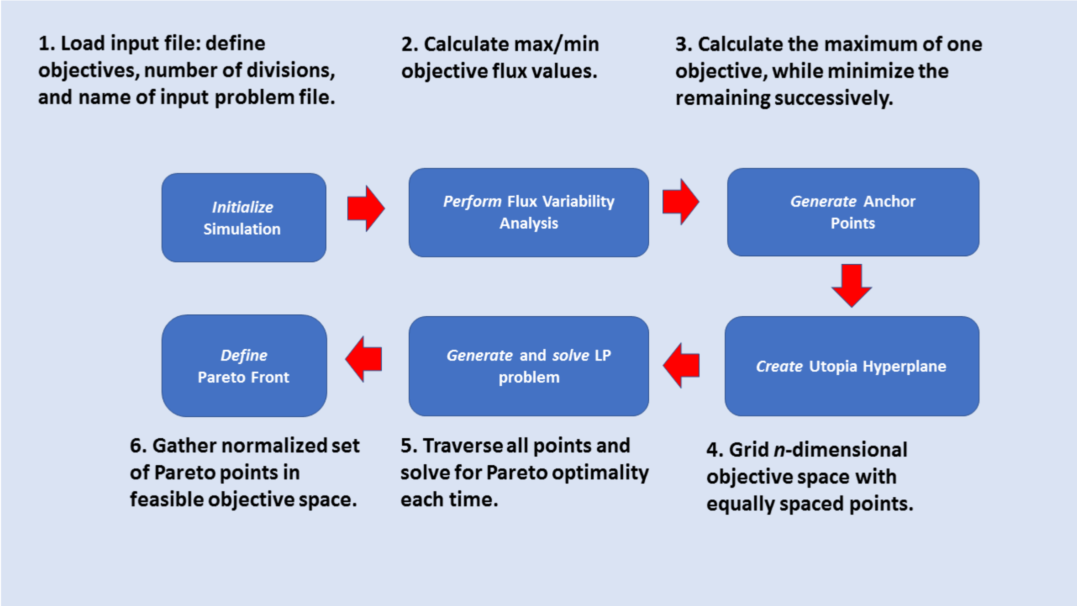

Figure M1: Schematic of MOFA workflow.

**Step 6: Gather Pareto points to form Pareto frontier.**
The set of Pareto points is combined into the frontier.

Step 5 is performed repeatly, traversing from point to point until all are covered. The simulation also skips points close to the previous point. Figure M1 summarizes and categorizes the workflow of the current version of the software.

### 7 Definitions

Flux variability analysis (FVA): for a given level of the cellular objective (e.g. biomass yield) finding the upper and lower bounds of all steady-state reaction fluxes can be determined.

Anchor point: Axis point in multi-objective space where the objective interest is at its maximum.

Utopia (hyper)plane: the multidimensional plane formed by the connection of the anchor points.

Pareto front(ier): The set of feasible points in objective spaceâĂŤpoints where moving away from it improves the value of the others.

COBRA Toolbox: Constraint-Based Reconstruction and Analysis add-on to MATLAB.

Normalized Normal Constraint (NNC) method: a multi-objective algorithm for generating an equally spaced set of Pareto points.

*Feasible: Feasible solution found.

*Wasted Simulation: Infeasible or unbounded solution found at point.

* Definition refers to variables in the code.
