## Supplementary material for "MOFA: Multi-Objective Flux Analysis for the COBRA Toolbox": Code files: anch_pts.m

```
%%%%%%%%%%%%%%%%%%%%%%%%%%%%%%%%%%%%%%%%%%%%%%%%%%%%%%%%%%%%
%                                                          %
%          MOFA(2017)                                      %
%          Marc Griesemer and Ali Navid                    %
%          Lawrence Livermore National Laboratory          %
%          Livermore, CA 94551 USA                         %
%                                                          %
%          function ap                                     %
%          input: model, model object                      %
%          input: mxhr, the vector of maximum fluxes       %
%          input: mihr, the vector of minimum fluxes       %
%          input: obar, the list of objectives             % 
%          input: obf, the main objective                  %
%          output: aphr, matrix of the anchor points       %
%                                                          %
%%%%%%%%%%%%%%%%%%%%%%%%%%%%%%%%%%%%%%%%%%%%%%%%%%%%%%%%%%%%

function [aphr] = anch_pts(model,mxhr, mihr, obar, obf)

objc = obar;
osize = length(objc);    
aphr = zeros(osize,osize);
bz = zeros(length(objc));
model.c(model.c==1) = 0;
%csense = cell(length(model.mets),1);
csense = [];
for i=1:length(model.mets)
   csense = [csense; 'S']; 
end
params=struct;
osense = -1;

     for i = 1:length(objc)

         for j=1:length(objc)
             
             if i ~= j
              model_temp = model;     
              % os2 = XX(j);
               model_temp.c(strcmp(model_temp.rxns,objc(j))==1) = 1;
               idx = find(strcmp(model_temp.rxns,objc(i))==1);
                
                  ub = mxhr(i);
                      
                  if abs(ub) <= 1e-6
                        ub = 0;
                  end
                 newm = ub - (1-.99999)*ub;
                 newx = ub + (0.00001)*ub;
                 

                  if ub >= 0
                    model_temp.ub(idx) = str2double(sprintf('%.5e',newx));
                    model_temp.lb(idx) = str2double(sprintf('%.5e',newm));
                 else
                    model_temp.lb(idx) = str2double(sprintf('%.5e',newx));
                    model_temp.ub(idx) = str2double(sprintf('%.5e',newm));  
                 end

                 % solve the LP problem for the current anch pt.
                 [x,faphr,origStat,extra] = glpk(model_temp.c,model_temp.S,model_temp.b,model_temp.lb,model_temp.ub,csense,[],osense,params);
                  
       
                 if (origStat == 180 || origStat == 5)
                      stat = 1; % Optimal solution found
                 elseif (origStat == 182 || origStat == 183 || origStat == 3 || origStat == 110)
                       stat = 0; % Infeasible
                 elseif (origStat == 184 || origStat == 6)
                      stat = 2; % Unbounded
                 else
                      stat = -1; % Solution not optimal or solver problem
                 end
                 
                 if stat == 1
                    mit = sprintf('%.5e',faphr);             
                 else
                    disp('Anchor points not calculated successfully.')
                    return;
                 end
            
                 aphr(i,j) = str2double(sprintf('%.3e',str2double(mit))); 
                 if abs(aphr(i,j)) < 1e-8
                    aphr(i,j) = 0.0; 
                 end
                 
             else
                 
                 aphr(i,j) = str2double(sprintf('%.3e',mxhr(i)));
                 if abs(aphr(i,j)) < 1e-8
                    aphr(i,j) = 0.0; 
                 end
             end
         end
     end
                 
end
```
