## Supplementary material for "MOFA: Multi-Objective Flux Analysis for the COBRA Toolbox": Code files: GPL.pdf

### The GNU General Public License (GPL)
### Version 2, June 1991

The following is the GPL open source license template to be used in each of the source files of a program where we assert LLNS copyright.

Attach the following notices to each source file. In this notice there should be a pointer to where the Full notice is to be found.

> Copyright (c) <YEAR>, Lawrence Livermore National Security, LLC. Produced at the Lawrence Livermore National Laboratory. Written by <code author's name and email>
>
> CODE-<R&R number> All rights reserved.
>
> This file is part of <code name>
>
> Please also read this link – Our Notice and GNU General Public License.
>
> This program is free software; you can redistribute it and/or modify it under the terms of the GNU General Public License (as published by the Free Software Foundation) version 2, dated June 1991.
>
> This program is distributed in the hope that it will be useful, but WITHOUT ANY WARRANTY; without even the IMPLIED WARRANTY OF MERCHANTABILITY or FITNESS FOR A PARTICULAR PURPOSE. See the terms and conditions of the GNU General Public License for more details.
>
> You should have received a copy of the GNU General Public License along with this program; if not, write to the Free Software Foundation, Inc., 59 Temple Place, Suite 330, Boston, MA 02111-1307 USA
>
> OUR NOTICE AND TERMS AND CONDITIONS OF THE GNU GENERAL PUBLIC LICENSE
>
> Our Preamble Notice
>
> A. This notice is required to be provided under our contract with the U.S. Department of Energy (DOE). This work was produced at the Lawrence Livermore National Laboratory under Contract No. DE-AC52-07NA27344 with the DOE.

B. Neither the United States Government nor Lawrence Livermore National Security, LLC nor any of their employees, makes any warranty, express or implied, or assumes any liability or responsibility for the accuracy, completeness, or usefulness of any information, apparatus, product, or process disclosed, or represents that its use would not infringe privately-owned rights.

C. Also, reference herein to any specific commercial products, process, or services by trade name, trademark, manufacturer or otherwise does not necessarily constitute or imply its endorsement, recommendation, or favoring by the United States Government or Lawrence Livermore National Security, LLC. The views and opinions of authors expressed herein do not necessarily state or reflect those of the United States Government or Lawrence Livermore National Security, LLC, and shall not be used for advertising or product endorsement purposes.

The precise terms and conditions for copying, distribution and modification follows.

GNU Terms and Conditions for Copying, Distribution, and Modification

0. This License applies to any program or other work which contains a notice placed by the copyright holder saying it may be distributed under the terms of this General Public License. The "Program," below, refers to any such program or work, and a "work based on the Program" means either the Program or any derivative work under copyright law: that is to say, a work containing the Program or a portion of it, either verbatim or with modifications and/or translated into another language. (Hereinafter, translation is included without limitation in the term "modification".) Each licensee is addressed as "you."

Activities other than copying, distribution and modification are not covered by this License; they are outside its scope. The act of running the Program is not restricted, and the output from the Program is covered only if its contents constitute a work based on the Program (independent of having been made by running the Program). Whether that is true depends on what the Program does.

1. You may copy and distribute verbatim copies of the Program's source code as you receive it, in any medium, provided that you conspicuously and appropriately publish on each copy an appropriate copyright notice and disclaimer of warranty; keep intact all the notices that refer to this License and to the absence of any warranty; and give any other recipients of the Program a copy of this License along with the Program.

You may charge a fee for the physical act of transferring a copy, and you may at your option offer warranty protection in exchange for a fee.

2 You may modify your copy or copies of the Program or any portion of it, thus forming a work based on the Program, and copy and distribute such modifications or work under the terms of Section 1 above, provided that you also meet all of these conditions:

a) You must cause the modified files to carry prominent notices stating that you changed the files and the date of any change.

b) You must cause any work that you distribute or publish, that in whole or in part contains or is derived from the Program or any part thereof, to be licensed as a whole at no charge to all third parties under the terms of this License.

c) If the modified program normally reads commands interactively when run, you must cause it, when started running for such interactive use in the most ordinary way, to print or display an announcement including an appropriate copyright notice and a notice that there is no warranty (or else, saying that you provide a warranty) and that users may redistribute the program under these conditions, and telling the user how to view a copy of this License. (Exception: if the Program itself is interactive but does not normally print such an announcement, your work based on the Program is not required to print an announcement.)

These requirements apply to the modified work as a whole. If identifiable sections of that work are not derived from the Program, and can be reasonably considered independent and separate works in themselves, then this License, and its terms, do not apply to those sections when you distribute them as separate work. But when you distribute the same section as part of a whole which is a work based on the Program, the distribution of the whole must be on the terms of this License, whose permissions for other licensees extend to the entire whole, and thus to each and every part regardless of who wrote it.

Thus, it is not the intent of this section to claim rights or contest your rights to work written entirely by you; rather, the intent is to exercise the right to control the distribution of derivative or collective works based on the Program.

In addition, mere aggregation of another work not based on the Program with the Program (or with a work based on the Program) on a volume of a storage or distribution medium does not bring the other work under the scope of this License.

3. You may copy and distribute the Program (or a work based on it, under Section 2) in object code or executable form under the terms of Sections 1 and 2 above provided that you also do one of the following:

a) Accompany it with the complete corresponding machine-readable source code, which must be distributed under the terms of Sections 1 and 2 above on a medium customarily used for software interchange; or,

b) Accompany it with a written offer, valid for at least three years, to give any third party, for a charge no more than your cost of physically performing source distribution, a complete machine-readable copy of the corresponding source code, to be distributed under the terms of Sections 1 and 2 above on a medium customarily used for software interchange; or,

c) Accompany it with the information you received as to the offer to distribute corresponding source code. (This alternative is allowed only for noncommercial distribution and only if you received the program in object code or executable form with such an offer, in accord with Subsection b above.)

The source code for a work means the preferred form the work for making modifications to it. For an executable work, complete source code means all the source code for all modules it contains, plus any associated interface definition files, plus the scripts used to control compilation and installation of the executable. However, as a special exception, the source code distributed need not include anything that is normally distributed (in either source or binary form) with the major components (compiler, kernel, and so on) of the operating system on which the executable runs, unless that component itself accompanies the executable.

If distribution of executable or object code is made by offering access to copy from a designated place, then offering equivalent access to copy the source code from the same place counts as distribution of the source code, even though third parties are not compelled to copy the source along with the object code.

4. You may not copy, modify, sublicense, or distribute the Program except as expressly provided under this License. Any attempt otherwise to copy, modify, sublicense or distribute the

Program is void, and will automatically terminate your rights under this License. However, parties who have received copies, or rights, from you under this License will not have their licenses terminated so long as such parties remain in full compliance.

2 You are not required to accept this License, since you have not signed it. However, nothing else grants you permission to modify or distribute the Program or its derivative works. These actions are prohibited by law if you do not accept this License. Therefore, by modifying or distributing the Program (or any work based on the Program), you indicate your acceptance of this License to do so, and all its terms and conditions for copying, distributing or modifying the Program or works based on it.

3 Each time you redistribute the Program (or any work based on the Program), the recipient automatically receives a license from the original licensor to copy, distribute or modify the Program subject to these terms and conditions. You may not impose any further restrictions on the recipients' exercise of the rights granted herein. You are not responsible for enforcing compliance by third parties to this License.

4 If, as a consequence of a court judgment or allegation of patent infringement or for any other reason (not limited to patent issues), conditions are imposed on you (whether by court order, agreement or otherwise) that contradict the conditions of this License, they do not excuse you from the conditions of this License. If you cannot distribute so as to satisfy simultaneously your obligations under this License and any other pertinent obligations, then as a consequence you may not distribute the Program at all. For example, if a patent license would not permit royalty-free redistribution of the Program by all those who receive copies directly or indirectly through you, then the only way you could satisfy both it and this License would be to refrain entirely from distribution of the Program.

If any portion of this section is held invalid or unenforceable under any particular circumstance, the balance of the section is intended to apply and the section as a whole is intended to apply in other circumstances.

It is not the purpose to this section to induce you to infringe any patents or other property right claims or to contest validity of any such claims; this section has the sole purpose of protecting the integrity of the free software distribution system, which is implemented by public license practices. Many people have made generous contributions to the wide range of software distributed through that system in reliance on consistent application of that system; it is up to the author/donor to decide if he or she is willing to distribute software through any other system and a licensee cannot impose that choice.

This section is intended to make thoroughly clear what is believed to be a consequence of the rest of this License.

1 If the distribution and/or use of the Program is restricted in certain countries either by patents or by copyrighted interfaces, the original copyright holder who places the Program under this License may add an explicit geographical distribution limitation excluding those countries, so that distribution is permitted only in or among countries not thus excluded. In such case, this License incorporates the limitation as if written in the body of this License.

9. The Free Software Foundation may publish revised and/or new versions of the General Public License from time to time. Such new versions will be similar in spirit to the present version, but may differ in detail to address new problems or concerns.

Each version is given a distinguishing version number. If the Program specifies a version number of this License which applies to it and "any later version," you have the option of following the terms and conditions either of that version of any later version published by the Free Software Foundation. If the Program does not specify a version number of this License, you may choose any version ever published by the Free Software Foundation.

2 If you wish to incorporate parts of the Program into other free programs whose distribution conditions are different, write to the author to ask for permission. For software which is copyrighted by the Free Software Foundation, write to the Free Software Foundation; we sometimes make exceptions for this. Our decision to grant permission will be guided by the two goals of preserving the free status of all derivatives of our free software and or promoting the sharing and reuse of software generally.

NO WARRANTY

11. BECAUSE THE PROGRAM IS LICENSED FREE OF CHARGE, THERE IS NO WARRANTY FOR THE PROGRAM, TO THE EXTENT PERMITTED BY APPLICABLE LAW. EXCEPT WHEN OTHERWISE STATED IN WRITING THE COPYRIGHT HOLDERS AND/OR OTHER PARTIES PROVIDE THE PROGRAM "AS IS" WITHOUT WARRANTY OF ANY KIND, EITHER EXPRESSED OR IMPLIED, INCLUDING, BUT NOT LIMITED TO, THE IMPLIED WARRANTIES OF MERCHANTABILITY AND

FITNESS FOR A PARTICULAR PURPOSE. THE ENTIRE RISK AS TO THE QUALITY AND PERFORMANCE OF THE PROGRAM IS WITH YOU. SHOULD THE PROGRAM PROVE DEFECTIVE, YOU ASSUME THE COST OF ALL NECESSARY SERVICING, REPAIR OR CORRECTION.

2 IN NO EVENT UNLESS REQUIRED BY APPLICABLE LAW OR AGREED TO IN WRITING WILL ANY COPYRIGHT HOLDER, OR ANY OTHER PARTY WHO MAY MODIFY AND/OR REDISTRIBUTE THE PROGRAM AS PERMITTED ABOVE, BE LIABLE TO YOU FOR DAMAGES, INCLUDING ANY GENERAL, SPECIAL INCIDENTAL OR CONSEQUENTIAL DAMAGES ARISING OUT OF THE USE OR INABILITY TO USE THE PROGRAM (INCLUDING BUT NOT LIMITED TO LOSS OF DATA OR DATA BEING RENDERED INACCURATE OR LOSSES SUSTAINED BY YOU OR THIRD PARTIES OR A FAILURE OF THE PROGRAM TO OPERATE WITH ANY OTHER PROGRAMS), EVEN IF SUCH HOLDER OR OTHER PARTY HAS BEEN ADVISED OF THE POSSIBILITY OF SUCH DAMAGES.

END OF TERMS AND CONDITIONS
