## Supplementary material for "MOFA: Multi-Objective Flux Analysis for the COBRA Toolbox": Code files: max_min.m

```
%%%%%%%%%%%%%%%%%%%%%%%%%%%%%%%%%%%%%%%%%%%%%%%%%%%%%%%%%%%%
%                                                          %
%          MOFA(2017)                                      %
%          Marc Griesemer and Ali Navid                    %
%          Lawrence Livermore National Laboratory          %
%          Livermore, CA 94551 USA                         %
%                                                          %
%          Function: max_min                               %
%          input: model, COBRA model object                %
%          input: objc, the full list of objectives        %
%          output: vector of maximum fluxes for objectives %
%          output: vector of maximum fluxes for objectives %
%                                                          %
%%%%%%%%%%%%%%%%%%%%%%%%%%%%%%%%%%%%%%%%%%%%%%%%%%%%%%%%%%%%

function [mxhr,mihr] = max_min(model,objc)

model.c(model.c==1) = 0;
mihr = zeros(length(objc),1);
mxhr = zeros(length(objc),1);

csense = [];
for ij=1:length(model.mets)
    
   csense = [csense; 'S']; 
end
length(csense)
% loop through the objectives
for i=1:length(objc)
    
    
    model_temp = model;
    obj = objc(i);
    model_temp.c(strcmp(model_temp.rxns,obj)==1) = 1;
    cc = model_temp.c;
    i
    
    % find the minimum flux for the ith objective
    objc(i)
    osense = 1;
    param = struct;
     % solve for the minimum flux  
    [xmin,fmin,origStat_min,extra_min] = glpk(cc,model_temp.S,model_temp.b,model_temp.lb,model_temp.ub,csense,[],osense,param);

     
    tmin = fmin;

    mit = sprintf('%.5e',tmin);
         
    mihr(i) = str2double(mit);
      
    % find the maximum flux for the ith objective
    osense = -1;
    
    % solve for the maximum flux
    [xmax,fmax,origStat_max,extra_max] = glpk(cc,model_temp.S,model_temp.b,model_temp.lb,model_temp.ub,csense,[],osense,param);  
     tmax = fmax;
     
     mxt = sprintf('%.5e',tmax);
   
     mxhr(i) = str2double(mxt);
                                      
end % end objc loop 
    for i=1:length(objc)
        if abs(mxhr(i)) < 1e-8
            mxhr(i) = 0.0;
        end
        if abs(mihr(i)) < 1e-8
           mihr(i) = 0.0; 
        end
        mxhr(i) 
        mihr(i)
    end
end %  function: max_min_sobe
```
