## Supplementary material for "MOFA: Multi-Objective Flux Analysis for the COBRA Toolbox": Code files: mofa.m

```
%%%%%%%%%%%%%%%%%%%%%%%%%%%%%%%%%%%%%%%%%%%%%%%%%%%%%%%%%%%%
%                                                          %
%  MOFA (2017)                                             %
%  Marc Griesemer and Ali Navid                            %
%  Lawrence Livermore National Laboratory                  %
%  Livermore, CA 94551 USA                                 %  
%                                                          %
% Inputs:                                                  % 
% model: COBRA model object                                %
% inp_file: the filename of the input file with the        % 
%  list of objectives                                      %
% obs: set of objectives: cell array of strings            %
% obf: main objective: cell containing 1 string            %
% div: number of divisions: number                         %
% mi_mx: min ('min') or max ('max'): string                %
%                                                          %
% Outputs:                                                 %
% mofa_sol: the Pareto front: cell array of strings        %
% mxhr: set of maximum fluxes for objectives, double       %
% mihr: set of minimum fluxes for objectives, double       %
% aphr: anchor points, 2D matrix of doubles                %
%                                                          %
%%%%%%%%%%%%%%%%%%%%%%%%%%%%%%%%%%%%%%%%%%%%%%%%%%%%%%%%%%%%

function [mofa_sol, mxhr, mihr, aphr] = mofa(model,inp_file,varargin)

mofa_sol = [];
mxhr = [];
mihr = [];
aphr = [];

% only glpk supported at this time, installed with COBRA Toolbox

status_glpk = changeCobraSolver('glpk');

if status_glpk == 0
    disp('Change to solver glpk failed')
    return; 
end

% checks number of input arguments
narginchk(2,6);

%%%%%%%%%%%%%%%%%%%%%%%%%%%%%%%%%%%%%%%%%%%%%%%
%  if user has input file of objectives
%  format (see User Manual or 'ecoli_input.txt)
%  ------------------------------
%  #comment
%  "main objective"
%  #comment
%  "other objectives" one-per-line
%  ------------------------------
%  reaction must be in COBRA model object
%  avoid blank lines at end/misspellings 
%%%%%%%%%%%%%%%%%%%%%%%%%%%%%%%%%%%%%%%%%%%%%%


%%%%%%%%%%%%%%%%%%%%%%%%%%%%%%%%%%%%%%%%%%%%%%%%%%%%%%%%%%%%%%%%%
%
% INPUT SECTION
%
%%%%%%%%%%%%%%%%%%%%%%%%%%%%%%%%%%%%%%%%%%%%%%%%%%%%%%%%%%%%%%%%%


% Empty or invalid model object
if isempty(fieldnames(model))
   disp('Invalid model object')
   return
end 


% testing for valid COBRA model structure
if ~isfield(model,'S')
    disp('Stoichiometric matrix (S) in model object missing or invalid');
    return
end
if ~isfield(model,'rxns')
    disp('Reactions (rxns) field in model object missing or invalid')
    return
end

if ~isfield(model,'mets')
    disp('Metabolites (mets) field in model object missing or invalid')
    return
end

if ~isfield(model,'lb')
    disp('Lower bounds (lb) field in model object missing or invalid')
    return 
end

if ~isfield(model,'ub')
    disp('Upper bounds (ub) field in model object missing or invalid')
    return
end

if ~isfield(model,'b') 
    disp('Flux vector (b) field in model object missing or invalid')
    return
end        


       
% check to see if using input file

if nargin > 2
  if isempty(inp_file) && isempty(varargin{1})
      disp('Objectives need to be defined in arguments or input file')
      disp('Exiting...')
      return;
  end    
end

if nargin > 3
  if isempty(inp_file) && (isempty(varargin{1}) || isempty(varargin{2}))
     disp('Objectives need to be defined in function arguments or input file')
     disp('Exiting...')
     return;
  end
end


% check for valid input file name and attempt to open
if ~isempty(inp_file)
 
 fn = inp_file;
 fid = fopen(fn,'r');
 
 if fid == -1
    Xy =  sprintf('Can''t open input file: %s', fn);
    disp(Xy)
    disp('Exiting...')
    return
 end
 
 INFO = [];

 % read the input file 
 % put the lines into a vector
 while ~feof(fid)
      
   c = cellstr(strtrim(fgetl(fid)));
   INFO = [INFO; c];
              
 end
 
 if length(INFO) <= 1
    disp('Error: empty input file')
    disp('Exiting...')
    return;
 end

% the main objective to optimize
 obf = INFO(2);
 obs = [];
 ii = 4;
 while ii < length(INFO)+1
   if ~strcmp(INFO(ii),'') 
    obs = [obs; INFO(ii)];
   end
   ii = ii+1;
 end
end

% unless objectives specified here, uses objectives from input file
if nargin > 2
    
     if ~isempty(varargin{1}) && ~isempty(varargin{2})
        obs = [];
        obf = [];
        obf = varargin{1};
        obs = reshape(varargin{2},length(varargin{2}),1)
         
     end
end

if isempty(obs) || isempty(obf)
     disp('Error: objectives not specified correctly. Use input file or function arguments.');
     disp('Exiting...')
     return;
end

if nargin < 5
   div = 10;
else
   div  = varargin{3};
   varargin{3}
end

if nargin < 6
   mi_mx = 'max'; 
else
   mi_mx = varargin{4};
   varargin{4}
end


if strcmp(mi_mx,'max');
   osense = -1; % default maximize
elseif strcmp(mi_mx,'min')
   osense = 1;
else
   disp('Error: argument ''mi_mx'' has to be ''min'' or ''max''')
   disp('Exiting...')
   return;
end

is_not_present1 = ~any(strcmp(model.rxns,obf));

if is_not_present1
    
   sprintf('Error: main objective ''%s'' is not a valid reaction in model.',obf{1})
   disp('Exiting...')
   return

end

obsSize = length(obs)

for i=1:obsSize
   is_not_present2 = ~any(strcmp(model.rxns,obs(i)));
   if is_not_present2
      sprintf('Error: objective ''%s'' is not a valid reaction in model.',obs{i})
      disp('Exiting...')
      return
   end

end

if div < 3
   disp('Minimum number of divisions is 3');
   div = 3;
end

tic

%%%%%%%%%%%%%%%%%%%%%%%%%%%%%%%%%%%%%%%%%%%%%%%%%%%%%%%%%%%%%%%%%
%
% INITIALIZATION SECTION
%
%%%%%%%%%%%%%%%%%%%%%%%%%%%%%%%%%%%%%%%%%%%%%%%%%%%%%%%%%%%%%%%%%


% initialize vectors as empty
hahr = [];
main_obj = [];
sol2 = [];

no_sim = 10000000; % large number for testing 

disp('Beginning MOFA calculation.');

% sort the objectives
sobs = sort(obs)

%%%%%%%%%%%%%%%%%%%%%%%%%%%%%%%%%%%%%%%%%%%%%%%%%%%%%%%%%%%

% copy the objectives 
obss = sobs;

%  add the main objective to optimize
sobss =    [obf; obss]; 

obssSize = length(sobss)

%%%%%%%%%%%%%%%%%%%%%%%%%%%%%%%%%%%%%%%%%%%%%%%%%%%%%%%%%%%%%%%%%

% find the maximum and minimum objective fluxes
disp('Calculating minimum and maximum objective fluxes...')
[mxhr,mihr] = max_min_sobe(model,sobss);

% for normalization correction, so no divide by 0
for i=1:obssSize
    if mihr(i) < 0 && mxhr(i) <= 0  
        mxhr(i) = abs(mihr(i));
    end
end

disp('Finished calculating minimum and maximum objective fluxes...')

%  Find the anchor points
aphr = zeros(length(sobss),length(sobss));

disp('Calculating anchor points...')
aphr =  anch_pts_sobe(model,mxhr,mihr,sobss,obf);
disp('Finished calculating anchor points...')


%%%%%%%%%%%%%%%%%%%%%%%%%%%%%%%%%%%%%%%%%%%%%%%%%%%%%%%%%%%%%%%%%

obv = aphr(1,1);
 
alpha = zeros(obssSize,1);
fxhr = zeros(obsSize,1);
 
% number of objectives minus the main one
 num = obsSize;
% 
% % initialize the alphas (points to iterate over)
for kd=1:num;
 
   fxhr(kd)  = mihr(kd+1);
   alpha(kd+1) = 0.0;
 
end

alpha(1) = 0.0;

% the size of a division
 division = 1.0/div;


% initialize the counters for total number of solves (sim) and
% infeasible/unbounded solutions (wastedsim)
% number of points    

count = 0;
wastedsim = 0;
sum = 0;
sim = 0;
feas = 0;

tol = 1e-3;

% set up the glpk problem parameters
csense = [];
for ij=1:length(model.mets)
    
   csense = [csense; 'S']; 
   
end % equality

% make output file for writing
MO = fopen('mofa_output.txt','w');
for i=2:length(sobss)
	fprintf(MO,'%s\t',char(sobss(i)));
end

fprintf(MO,'%s\n',char(sobss(1)));
fclose('all');
   
% the main loop to iterate and solve over all points
while alpha(2) <= 1.0 && count < no_sim 
    
    sum = 2.0;

    while sum > 1.0
 
       sum = 0.0;
       
       for ii=1:length(sobss)   
          if abs(alpha(ii)) < 1e-6
              alpha(ii) = 0.0;
          end
       end
       
       if alpha(2) > 1.0 
           
	       break
       end
       

       for n=0:num-1
                             
          sum = sum + alpha(num-n+1);
               
          if sum > 1.0
                  
             if alpha(num-n+1) > 1.0
                  
                alpha(num-n)  =  alpha(num-n) + division;
                alpha(num-n+1) = 0.0;

 		     else
 
                alpha(num) = alpha(num) + division;
 		        alpha(num+1) = 0.0;
                  
             end
                       
             break
             
          end 
        end   % for loop
    end       %  while(sum > 1.0) loop
    
 

    if sum <= 1 

         alpha(1) = 1.0 - sum;
    end
    
   
    if sum > 1

         continue
    end
    
    for i=1:obsSize
       
      fxhr(i) = 0.0;
       
          for j=1:obsSize+1
 
             fxhr(i) = fxhr(i) +  alpha(j)*aphr(j,i+1);

          end
          
          hahr = [hahr; fxhr(i)];
              
    end
    
% this section is to test whether one can skip over nearby values
test = 0;

 for k=0:count-1

        if k == count -1

          break

        end
  
 test = 0;
  
    for kk = 1:obsSize

      dx = fxhr(kk) -  hahr(k*obsSize+kk);
      % tol=1e-3 is the default tolerance for now
      if abs(dx) < tol*fxhr(kk)
      
            
               test = test +  1;

      else
               break
      end
    end  
   
    if test == num
 
        break

    end
end
 
if test == num
    
   alpha(num+1) = alpha(num+1) +  division; 
   

   count = count + 1
 
   continue
end
 
if alpha(2) > 1
 
   break
end
 
%%%%%%%%%%%%%%%%%%%%%%%%%%%%%%%%%%%%%%%%%%%%%%%%%%%%%%%%%%%%%%%%%
% 
%  this is where you solve the LP problem
% 
% 
%%%%%%%%%%%%%%%%%%%%%%%%%%%%%%%%%%%%%%%%%%%%%%%%%%%%%%%%%%%%%%%%%

model_temp  = model;

% calculate the objective values at this point
for kc=1:obsSize
     
      if abs(kc) < 1e-6
          
          fxhr(kc) = 0.0;
      
      end 
      % update bounds of objectives
      newm = fxhr(kc)*(0.999);
      newx = fxhr(kc)*(1.001);
      
      if newx > newm
                 
           model_temp.lb(strcmp(model_temp.rxns,char(sobs(kc)))==1) = newm;
           model_temp.ub(strcmp(model_temp.rxns,char(sobs(kc)))==1) = newx;
      else
           model_temp.lb(strcmp(model_temp.rxns,char(sobs(kc)))==1) = newx;
           model_temp.ub(strcmp(model_temp.rxns,char(sobs(kc)))==1) = newm;           
       
      end       
end
      
   
       cc = model_temp.c;
       A = model_temp.S;
       b = model_temp.b;
       lb = model_temp.lb;
       ub = model_temp.ub;
       params= struct; % empty placeholder struct, not used
       
       [x,f,origStat,extra] = glpk(cc,A,b,lb,ub,csense,[],osense,params);
       
       if (origStat == 180 || origStat == 5)
             stat = 1; % Optimal solution found
       elseif (origStat == 182 || origStat == 183 || origStat == 3 || origStat == 110)
             stat = 0; % Infeasible
       elseif (origStat == 184 || origStat == 6)
             stat = 2; % Unbounded
       else
             stat = -1; % Solution not optimal or solver problem
       end
   % infeasible/unbounded solutions
   if stat ~= 1
       
      % count as wasted simulation
 
      wastedsim  = wastedsim + 1;
   else 
  
    % feasible solutions
    
       v = f;
       
       % normalize the main objective value
       normval = abs(v)/obv;

       sol1= [];
      WO =  fopen('mofa_output.txt','a');
      for k=2:obsSize+1
          % normalize the remaining objective value

          if mxhr(k) ~= 0
             u(k-1) = abs(fxhr(k-1)/mxhr(k));
          else
             disp('Divide by zero...exiting!');
          end
          
          uuu = '';
          uuu = sprintf('%.2f',u(k-1));
          fprintf(WO,'%.2f\t',u(k-1));
          sol1= [sol1; str2double(uuu)];
         
             
      end
      
        % increment number of feasible solutions
         feas = feas + 1;

         normval = str2double(sprintf('%.2f',normval));
         fprintf(WO, '%.2f\n',normval);
         fclose('all');
         sol2 = [sol2; sol1'];
         main_obj = [main_obj; normval];
    end
     
   % move to the next point
   alpha(num+1) = alpha(num+1) +  division; 
   
   % advance simulation
   count = count + 1
   sim = sim + 1
   wastedsim
   feas

 end
 % construct the final solution

 mofa_sol = horzcat(sol2,main_obj);

 % total counts, feasible counts, wasted simulations at end
 
 % total iterations
 sprintf('Total count of iterations: %d\n',count);
  % total feasible points
 sprintf('Total feasible solutions: %d\n',feas);
 sprintf('Total wasted simulations: %d\n',wastedsim);


 toc
 end
```
